# MapCaller – An integrated and efficient tool for short-read mapping and variant calling using high-throughput sequenced data

**DOI:** 10.1101/783605

**Authors:** Hsin-Nan Lin, Wen-Lian Hsu

## Abstract

With the advance of next-generation sequencing (NGS) technologies, more and more medical and biological researches adopt NGS technologies to characterize the genetic variations between individuals. The identification of personal genome variants using NGS technology is a critical factor for the success of clinical genomics studies. It requires an accurate and consistent analysis procedure to distinguish functional or disease-associated variants from false discoveries due to sequencing errors or misalignments. In this study, we integrate the algorithms for read mapping and variant calling to develop an efficient and versatile NGS analysis tool, called MapCaller. It not only maps every short read onto a reference genome, but it also detects single nucleotide variants, indels, inversions and translocations at the same time. We evaluate the performance of MapCaller with existing variant calling pipelines using three simulated datasets and four real datasets. The result shows that MapCaller can identify variants accurately. Moreover, MapCaller runs much faster than existing methods. It is available at https://github.com/hsinnan75/MapCaller.

## Background

With the advance of next-generation sequencing (NGS) technologies, it is becoming affordable to support various applications of precision medicine in the near future^1^. More and more medical and biological researches adopt NGS technologies to characterize the genetic variations between individuals ^2,3^. Such genetic variations can be classified into three types: (1) single nucleotide variant (SNV, also referred to as SNP); (2) insertion and deletion (indel); and (3) structural variant (SV, including translocation, inversion, copy number variation and indels of size at least 50 bp).

The identification of genome variants is a critical factor for the success of clinical genomics studies^4^. It requires an accurate and consistent analysis procedure to distinguish true variants from false discoveries. This procedure often involves the steps of short read alignment, alignment rearrangement and variant calling. In each step, one or more tools are applied to generate desired output. For example, BWA^5^, Bowtie ^6,7^, GEM ^8^, Subread ^9^, HISAT/HISAT2 ^10^ and KART ^11^ are read aligners that can map NGS short reads onto a reference genome and generate their alignments. SAMtools ^12^ and Picard ^13^ provide various utilities for manipulating read alignments. For variant calling, the Genome Analysis Tool Kit (GATK) ^14^, Freebayes ^15^, Platypus ^16^, VarScan ^17^, DeepVariant^18^ and SAMtools are widely used. Different combinations of those tools produce various analysis pipelines. Different variant calling pipelines may generate substantial disagreements of variant calls. Several studies ^4,19^ have been conducted to confirm the disagreements of variant calling among different pipelines. Besides, all existing variant calling pipelines are time and space consuming. They require read alignments are sorted and stored in desired format.

In this study, we present MapCaller, an efficient and versatile NGS analysis tool, by integrating the algorithms for read mapping and variant calling. For read mapping, we adopt a divide-and-conquer strategy to separate a read into regions with and without gapped alignment. With this strategy of read partitioning, SNVs, indels and breakpoints can be identified efficiently. For variant calling, MapCaller maintains a position frequency matrix to keep track of every nucleobase’s occurrence at each position of the reference genome while mapping the read sequences. Since MapCaller collects all information required for variant identification while reads are mapped onto the reference genome, variants can be called directly in the same process. Therefore, the conventional analysis pipeline can be simplified greatly. Most existing variant callers can only detect a few specific types of variants, however MapCaller can detect multiple types of variations, including SNVs, indels, inversions and translocations. We demonstrate that MapCaller not only produces comparable performance on variant calling, but it also spends much less time compared to selected variant calling pipelines. MapCaller was developed under Linux 64-bit environment and implemented with standard C/C++. It takes read files (FASTA/FASTQ) as input and outputs all predicted variants in VCF format. The source codes of MapCaller and benchmark datasets are available at https://github.com/hsinnan75/MapCaller.

## Results and Discussion

### Experiment design

We develop a simulator to generate genome variations using the E.Coli K-12 strain, human chromosome 1 and whole human reference genome (GRCh38). The simulator (please refer to Supplementary data S4) randomly generates sequence variations with occurrences of 2700 substitutions, 180 small indels (1∼10 bp), 45 large indels (11∼50 bp), 1 translocation (TNL, size ranges 1000∼2000bp), 1 inversion (INV, size ranges 1000∼2000bp) and 1 copy number variation (CNV, size ranges 300∼1300bp) for every 1,000,000 base pairs. We use WGSIM (https://github.com/lh3/wgsim) to generate simulated short read sequences for each mutant genome. The simulated read coverage is 30X, sequencing error rate is 2% and read length is 100bp. The synthetic datasets are referred to as Sim_Ecoli, Sim_Chr1 and Sim_HG, respectively. We also download four real NGS datasets from SRA web site, two from sample of HG001-NA12878 (RUN: SRR6062143 and SRR7781445) and two from sample of HG002-NA24385 (RUN: SRR3440404 and SRR6691661), where the dataset of SRR3440404 includes short reads from RUNs of SRR3440404 to SRR3440422 in order to have enough read depth. Note SRR6062143 and SRR7781445 are two separate runs of Illumina sequencing data of NA12878. SRR3440404 and SRR6691661 are also two separate runs of NA24385. GIAB (The Genome in a Bottle Consortium) ^20,21^ provides high-confidence SNP, small indel calls for the two sample genomes. They can be found at ftp://ftp-trace.ncbi.nlm.nih.gov/giab/ftp/release/. Those variant calls were made by GATK, Freebayes and Sentieon. Table 1 shows the number of short reads of each dataset as well as the number of each type of variants. It is noteworthy that GIAB does not provide structural variant annotation for samples NA12878 and NA24385.

**Table 1.** The benchmark datasets. Sim_Ecoli, Sim_Chr1, and Sim_HG are synthetic datasets. SRR6062143 and SRR7781445 are generated from NA12878 (HG001); SRR3440404 and SRR6691661 are generated from NA24385 (HG002). TNL: translocation; INV: inversion.

| Dataset | # of short reads | read length | SNV | indels | TNL | INV |
| --- | --- | --- | --- | --- | --- | --- |
| Sim_Ecoli | 1,472,676 | 100 | 12,718 | 1,109 | 2 | 4 |
| Sim_Chr1 | 78,859,696 | 100 | 627,576 | 52,067 | 261 | 211 |
| Sim_HG | 978,158,892 | 100 | 8,003,103 | 666,213 | 2,650 | 2,559 |
| SRR6062143 | 1,558,904,052 | 101 | 3,084,732 | 534,739 | NA | NA |
| SRR7781445 | 609,362,230 | 151 | 3,084,732 | 534,739 | NA | NA |
| SRR3440404 | 487,042,582 | 250 | 3,076,552 | 519,569 | NA | NA |
| SRR6691661 | 638,722,972 | 151 | 3,076,552 | 519,569 | NA | NA |

A conventional analysis pipeline includes a read mapper, SAM/BAM file processing and a variant caller. Thus, in study, we compare the performance of MapCaller with different combinations of those read mappers and variant callers. For read mapping, we select KART, BWA-MEM (BWA for short), Bowtie2 and GEM. For the variant calling, we select Genome Analysis Tool Kit HaplotypeCaller (GATK for short), Freebayes, SAMtools mpileup (Mpileup for short), DeepVariant and Platypus. For SAM/BAM file processing, we use SAMtools view/sort to perform file format converting and alignment sorting. We also compare the performance of structural variant calling with existing methods, including DELLY^22^, LUMPY^23^ and SVDetect^24^. The commands as well as the argument setting used for each pipeline are shown in Supplementary material (S3).

It is noteworthy that since MapCaller integrates the algorithms of read mapping and variant calling, it handles the whole procedure of analysis pipeline alone. The run time is estimated from the read mapping to variant calling. We estimate the precision, sensitivity and F1 score for each dataset using the RTG tools^25^. RGT’s vcfeval compares two variant sets and aims to maximize true positives and minimize false positives by finding the optimal variant score threshold. The optimal threshold achieves the TP/FP balance. We followed the same evaluation metric^18^ to allow errors in the determination of the exact variant alleles and genotypes. Thus, a true positive is a called variant that matches a reference variant, and a false positive is one that does not match any reference variant; Likewise, a false negative is a reference variant that does not match any called variant. Based on the number of true positives (TP), false positives (FP) and false negatives (FN), we define the precision, sensitivity (recall) and F1 score as precision = TP / (TP + FP), sensitivity = TP / (TP + FN) and F1 = 2×precision×sensitivity/ (precision + sensitivity).

### Performance comparison on synthetic datasets

We test every combination of read mapper and variant caller. We find that BWA-MEM combined with any selected variant caller generally performs better than any other selected read mapper; therefore, we only show the performance of pipelines involved with BWA-MEM here to compare to MapCaller. Table 2 summarizes the comparison result on the three synthetic datasets. The detailed comparison result can be found in the Supplementary data (S5).

**Table 2.**
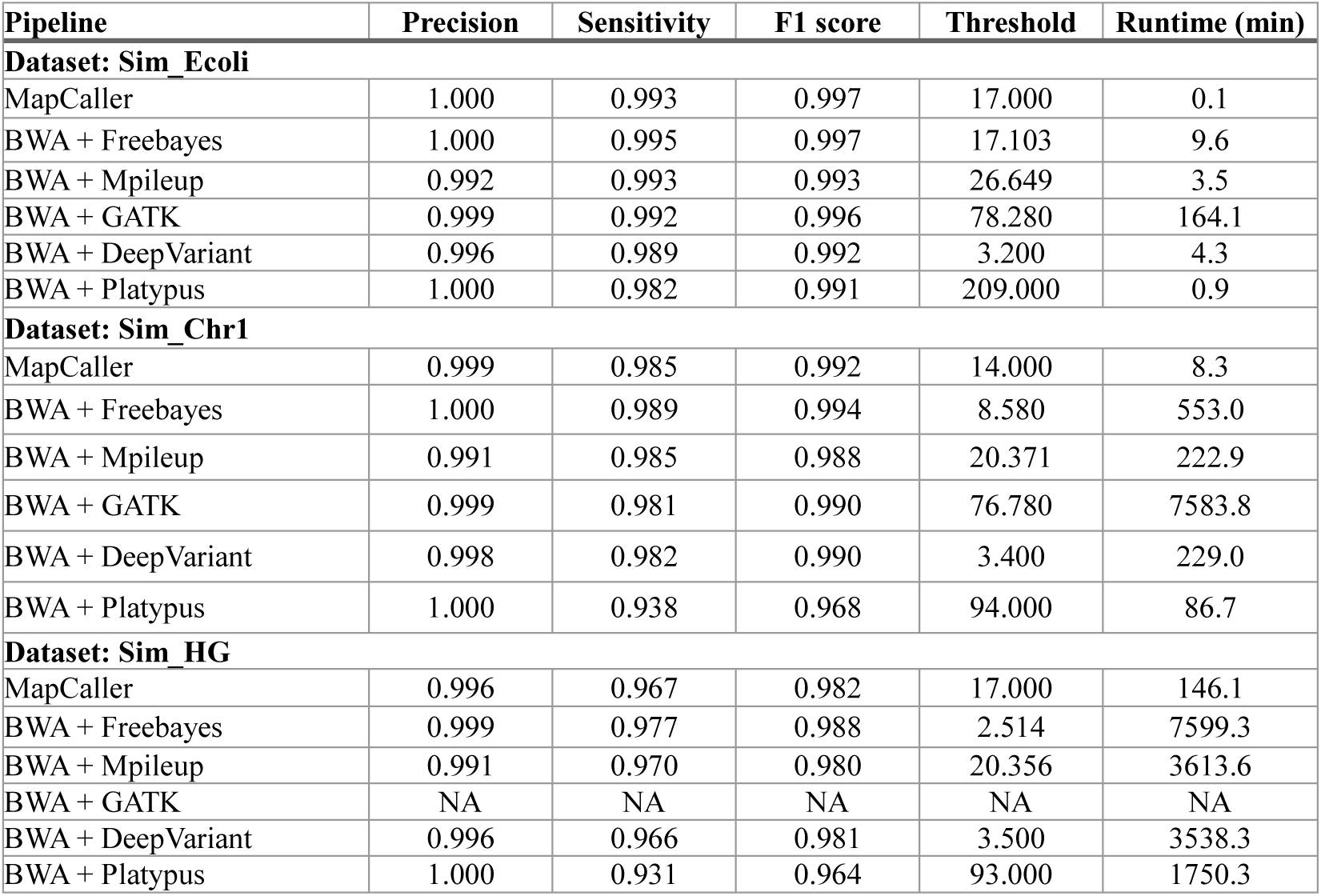
The performance comparison on the three synthetic datasets. We used the RTG tools to evaluate the performance of each variant calling method. RTG’s vcfeval compares two VCF files and selects the optimal QUAL threshold to maximize the F1-score.

| Pipeline | Precision | Sensitivity | F1 score | Threshold | Runtime (min) |
| --- | --- | --- | --- | --- | --- |
| **Dataset: Sim_Ecoli** |  |  |  |  |  |
| MapCaller | 1.000 | 0.993 | 0.997 | 17.000 | 0.1 |
| BWA + Freebayes | 1.000 | 0.995 | 0.997 | 17.103 | 9.6 |
| BWA + Mpileup | 0.992 | 0.993 | 0.993 | 26.649 | 3.5 |
| BWA + GATK | 0.999 | 0.992 | 0.996 | 78.280 | 164.1 |
| BWA + DeepVariant | 0.996 | 0.989 | 0.992 | 3.200 | 4.3 |
| BWA + Platypus | 1.000 | 0.982 | 0.991 | 209.000 | 0.9 |
| **Dataset: Sim_Chr1** |  |  |  |  |  |
| MapCaller | 0.999 | 0.985 | 0.992 | 14.000 | 8.3 |
| BWA + Freebayes | 1.000 | 0.989 | 0.994 | 8.580 | 553.0 |
| BWA + Mpileup | 0.991 | 0.985 | 0.988 | 20.371 | 222.9 |
| BWA + GATK | 0.999 | 0.981 | 0.990 | 76.780 | 7583.8 |
| BWA + DeepVariant | 0.998 | 0.982 | 0.990 | 3.400 | 229.0 |
| BWA + Platypus | 1.000 | 0.938 | 0.968 | 94.000 | 86.7 |
| **Dataset: Sim_HG** |  |  |  |  |  |
| MapCaller | 0.996 | 0.967 | 0.982 | 17.000 | 146.1 |
| BWA + Freebayes | 0.999 | 0.977 | 0.988 | 2.514 | 7599.3 |
| BWA + Mpileup | 0.991 | 0.970 | 0.980 | 20.356 | 3613.6 |
| BWA + GATK | NA | NA | NA | NA | NA |
| BWA + DeepVariant | 0.996 | 0.966 | 0.981 | 3.500 | 3538.3 |
| BWA + Platypus | 1.000 | 0.931 | 0.964 | 93.000 | 1750.3 |

It can be observed that MapCaller and the other selected pipelines perform comparably. They produce similar performance on the three synthetic datasets. Their F1 scores are around 0.96 ∼ 0.99. It implies that those methods are equally accurate and sensitive. For example, the precision, sensitivity and F1 score of MapCaller on Sim_HG are 0.996, 0.967 and 0.982, respectively. When further analyze some of the false negative cases, we find that most of false negatives occur at repetitive or highly similar regions. It is noteworthy that GATK spends too much time on Sim_HG and it fails to finish the whole process for all chromosomes within two weeks. However, it produces highly accurate results on Sim_Ecoli and Sim_Chr1.

It can be also observed that MapCaller is the fastest method among the selected methods. It only spends 146.1 minutes on Sim_HG, while the second fastest pipeline, BWA+Platypus spends 1750.3 minutes. It implies that MapCaller is at least 10 time faster than those selected pipelines. It is also noteworthy that the QUAL threshold of MapCaller, Mpileup, GATK and DeepVariant is more consistent throughout the three synthetic datasets. In contrast with the former methods, Freebayes and Platypus produce the best F1 scores at very different threshold. For example, the thresholds of Freebayes on the three datasets are 17.103, 8.580 and 2.514, respectively. It indicates that it is more challenging to determine an appropriate threshold if there are no reference calls for comparison.

### Performance comparison on real datasets

We download four real NGS datasets sequenced for genomes of NA12878 (HG001) and NA24385 (HG002). However, the two genomes do not have ground truth variant annotation. GIAB provides reference calls that were made by GATK, Freebayes and Sentieon for the sample genomes using datasets from at most five sequencing platforms (including Illumina, CG, Ion, 10X and Solid). In this study, we use the reference calls as the gold standard to estimate the performance of each method. We download two separate datasets from each sample genome because we would like to analyze if a variant caller would produce different results on different datasets from the same sample genome.

We summarize the comparison result in Table 3. It can be observed that DeepVariant performs the best among the selected methods. It’s F1 scores are all around 0.88∼0.89. It is not surprising that DeepVariant outperforms the other methods on the four real datasets since DeepVariant was trained with NA12878 datasets. Moreover, it is noteworthy that NA12878 and NA24385 are two different sample genomes, but they share around 60% variants. Platypus’s algorithms involve a sophisticated likelihood estimation model to estimate the population haplotype frequencies. It produces the best precisions among those selected methods on the four real datasets. However, it sacrifices the sensitivities slightly. MapCaller, GATK, Freebayes and Mpileup produce comparable performance. Their F1 scores are around 0.83∼0.86 for the four datasets.

**Table 3.** The performance comparison on the four real datasets. We used the RTG tools to evaluate the performance of each variant calling method. RTG’s vcfeval compares two VCF files and selects the optimal QUAL threshold to maximize the F1-score.

| Pipeline | Precision | Sensitivity | F1 score | Threshold | Runtime (hour) |
| --- | --- | --- | --- | --- | --- |
| **Dataset: SRR6062143 (NA12878)** |  |  |  |  |  |
| MapCaller | 0.771 | 0.941 | 0.848 | 12.000 | 2.0 |
| BWA + Freebayes | 0.793 | 0.962 | 0.869 | 147.135 | 122.7 |
| BWA + Mpileup | 0.787 | 0.919 | 0.848 | 28.422 | 72.1 |
| BWA + GATK | 0.784 | 0.945 | 0.857 | 269.520 | 203.8 |
| BWA + DeepVariant | 0.824 | 0.973 | 0.892 | 21.900 | 46.8 |
| BWA + Platypus | 0.829 | 0.917 | 0.871 | 132.000 | 25.6 |
| **Dataset: SRR7781445 (NA12878)** |  |  |  |  |  |
| MapCaller | 0.771 | 0.926 | 0.842 | 10.000 | 1.5 |
| BWA + Freebayes | 0.771 | 0.963 | 0.856 | 44.375 | 122.3 |
| BWA + Mpileup | 0.787 | 0.909 | 0.844 | 38.469 | 68.3 |
| BWA + GATK | 0.756 | 0.952 | 0.843 | 87.730 | 161.8 |
| BWA + DeepVariant | 0.814 | 0.972 | 0.886 | 31.200 | 62.3 |
| BWA + Platypus | 0.820 | 0.906 | 0.861 | 21.000 | 41..6 |
| **Dataset: SRR3440404 (NA24385)** |  |  |  |  |  |
| MapCaller | 0.757 | 0.931 | 0.835 | 10.000 | 2.1 |
| BWA + Freebayes | 0.763 | 0.968 | 0.853 | 128.672 | 116.4 |
| BWA + Mpileup | 0.795 | 0.904 | 0.846 | 77.000 | 59.9 |
| BWA + GATK | 0.779 | 0.956 | 0.858 | 152.730 | 227.9 |
| BWA + DeepVariant | 0.808 | 0.969 | 0.881 | 27.100 | 61.4 |
| BWA + Platypus | 0.800 | 0.910 | 0.852 | 67.000 | 25.3 |
| **Dataset: SRR6691661 (NA24385)** |  |  |  |  |  |
| MapCaller | 0.762 | 0.934 | 0.839 | 10.000 | 1.6 |
| BWA + Freebayes | 0.762 | 0.967 | 0.852 | 93.526 | 96.5 |
| BWA + Mpileup | 0.789 | 0.909 | 0.845 | 62.000 | 47.8 |
| BWA + GATK | 0.774 | 0.973 | 0.862 | 128.730 | 157.1 |
| BWA + DeepVariant | 0.807 | 0.969 | 0.880 | 26.200 | 44.5 |
| BWA + Platypus | 0.812 | 0.887 | 0.848 | 48.000 | 20.0 |

We further analyze MapCaller’s false negatives on SRR6062143. We find that there are 93948, 374289, 954752, 2175218 and 21264 reference variants are called from one, two, three, four and five platforms, respectively. However, the false negative rates are 0.375, 0.234, 0.061, 0.008 and 0.01 with respective to the number of platforms. It implies that MapCaller produces much less false negative rates for the variants from a greater number of platforms. Moreover, some of false negative cases are due to poor alignment quality and their read alignments are discarded by MapCaller. For example, the reference calls report three adjacent variants, which are chr1:20761541 (an insertion: AGAG), chr1:20761544 (SNV: C) and chr1:20761545 (a deletion: AT). Since there are more than three state transitions in the corresponding alignment, MapCaller generate the corresponding alignments and then they are considered poor alignment and discarded. If the adjacent variants appear in two separated alignments, they still can be called by MapCaller. It is estimated NA12878 contains 11,018 indel events (2.1%) that are adjacent to one another within five nucleotides, and NA24385 contains 10016 variants (2.0%). By contrast, MapCaller produces around 0.7% of indels that are adjacent within five nucleotides in average.

MapCaller is still much faster than any other pipeline on the real datasets. It is around 100 times faster than BWA+GATK. It spends around one or two hours to handle a human genome data with around 30X of read depth. If we only consider the run time for variant calling, MapCaller is much faster than GATK. For example, MapCaller spends three minutes on variant calling for SRR6062143, while GATK spends 203.8 hours for the same dataset. Though MapCaller runs very fast, it produces comparable result as Freebayes and GATK do in this analysis. Thus, we demonstrate MapCaller is a highly efficient variant calling method. Platypus is the second fastest method. It is the only method that can identify variants with multiple threads. That’s why it saves a lot of run time.

### Performance comparison on structural variant detection

In addition to SNVs and indels, MapCaller is also capable of identifying structural variants (referred to as SVs in the following description) simultaneously. We do not identify CNVs in this study. Table 4 shows the performance comparison on SV detection for MapCaller and other selected callers. We estimate the precision and recall of each caller. Variant called by SVDetect with “nb_pairs” < 50 are discarded. It can be observed that MapCaller produces high precisions and recalls on all the three simulated datasets. In particular, MapCaller produces the highest precisions among these callers. LUMPY produces high precisions and recalls on inversion detections. However, it produces relatively lower precisions and recalls on translocation detections. SVDetect produces high precision and recalls both on translocation and inversion detection. Similar to LUMPY, DELLY generally produces high precisions and recalls on inversion detections, but it cannot detect any translocation events on the three datasets. In summary, MapCaller and SVDetect can handle both translocation and inversion detection at high precision and recall.

**Table 4.** Performance comparison on structural variant detection.

| Data set | Method | Translocation |  |  | Inversion |  |  |
| --- | --- | --- | --- | --- | --- | --- | --- |
|  |  | Precision | Recall | F1 score | Precision | Recall | F1 score |
| Sim_Ecoli | MapCaller | 1.000 | 1.000 | 1.000 | 1.00 | 1.000 | 1.000 |
|  | LUMPY | 1.000 | 0.667 | 0.800 | 1.000 | 1.000 | 1.000 |
|  | SVDetect | 1.000 | 1.000 | 1.000 | 1.000 | 1.000 | 1.000 |
|  | DELLY | NA | NA | NA | 1.000 | 1.000 | 1.000 |
| Sim_Chr1 | MapCaller | 1.000 | 0.943 | 0.971 | 1.000 | 0.877 | 0.935 |
|  | LUMPY | 0.842 | 0.333 | 0.477 | 0.995 | 0.929 | 0.961 |
|  | SVDetect | 0.941 | 0.954 | 0.948 | 0.949 | 0.901 | 0.924 |
|  | DELLY | NA | NA | NA | 0.969 | 0.957 | 0.963 |
| Sim_HG | MapCaller | 0.999 | 0.948 | 0.973 | 1.000 | 0.878 | 0.935 |
|  | LUMPY | 0.879 | 0.353 | 0.504 | 1.000 | 0.937 | 0.968 |
|  | SVDetect | 0.874 | 0.930 | 0.901 | 0.968 | 0.891 | 0.928 |
|  | DELLY | NA | NA | NA | 0.991 | 0.952 | 0.971 |

## Conclusion

In this manuscript, we present MapCaller, an integrated system for read mapping and variant calling. MapCaller collects read alignment information during read mapping and maintains a position frequency matrix to keep track of alignments at each position of the reference genome. We evaluate the performance of MapCaller and the selected variant calling pipelines using three synthetic datasets and four real data sets from human genomes. The comparison results show that MapCaller not only performs comparably with existing methods, but it also spends the least amount of time. MapCaller is also versatile. It is capable of identifying SNVs, INDELs and structural variants simultaneously.

Since more and more medical and biological researches adopt NGS technologies to characterize the genetic variations between individuals, we believe MapCaller is able to provide accurate and reliable variant calls much faster than existing methods.

## Methods

MapCaller aligns every short read onto a reference genome and collects the alignment information during read mapping to identify sequence variants. MapCaller uses a modified algorithm of KART to perform read mapping. It maintains a position frequency matrix to keep track of every nucleobase’s frequency at each position in the reference genome and collects all insertion and deletion events that are found during read mapping. Furthermore, it gathers all possible breakpoints from discordant or partial read alignments. Finally, MapCaller finds sequence variants based on all of the above-mentioned information. The novelty of our algorithm derives from the integration of read mapping and variation information gathering into a coherent system for variant calling.

### Read mapping and alignment profiles

The details of read mapping method, KART can be found in our previous study ^11^. Here we focus on the high-level methodology description. KART adopts a divide-and-conquer strategy to handle matches and mismatches separately between read sequence and reference genome. KART identifies all locally maximal exact matches (LMEMs). We then cluster them according to their coordinates and fill gaps between LMEMs to create final alignments. In doing so, KART divides a read alignment into two groups: simple region pairs (abbreviated as *simple pairs*) and normal region pairs (*normal pairs*), where all simple pairs are LMEMs and normal pairs are gaps between simple pairs and might require gapped alignment (due to mismatches or indels). Thus, all SNVs and indels should be found in normal pairs, and breakpoints can be detected from normal pairs at either end of read alignment. We will explain the identification metrics of each variant type below.

For the convenience of describing the methodology of MapCaller, we define the following notations. Given a read sequence *R*, the reference genome *G*, Let *R*_*i*_ be the *i*-th residue of *R* and *R*[*i*_1_, *i*_2_] be the substring between *R*_*i*1_ and *R*_*i*2_. Likewise, let *G*_*j*_ be the *j*-th nucleotide of *G* and *G*[*j*_1_, *j*_2_] be the substring between *G*_*j*1_ and *G*_*j*2_. A simple pair (or a normal pair) consists of a read’s substring and its counterpart of reference’s substring. They can be represented as (*R*[*i*_1_, *i*_2_], *G*[*j*_1_, *j*_2_]). One or more simple/normal pair forms a candidate alignment. We perform pairwise alignment for each normal pair in a candidate alignment.

We check the alignment quality of a normal pair at either end of the read sequence to determine whether they should be discarded. The quality evaluation is as follows. Given an alignment of a normal pair at either end, MapCaller counts the number of mismatches and state transitions of the alignment. A state transition is an alignment state change of base alignment, insertion and deletion. If there are more than three state transitions or the number of mismatches is more than 0.3*N* (*N* is the number of base alignments) in the normal pair, it will be discarded from the candidate alignment. Such normal pair may appear due to false mapping or structural variants. MapCaller infers breakpoints based on such normal pairs.

Fig. 1 illustrates two cases of read alignments that reveal a translocation event and an inversion event, respectively. In Fig. 1(A), paired-end reads Read1 and Read2 are mapped far away from each other (could be mapped to different chromosomes) due to a translocation event in the sample genome. Thus, the two alignments are considered discordant. Read1 and Read2 are clipped a few bases since they cover a breakpoint. Likewise, in Fig. 1(B), paired-end reads Read3 and Read4 are mapped with same orientation due to an inversion event in the sample genome. The two alignments are also considered discordant. If a read covers a breakpoint, its alignment will be clipped by the breakpoint. MapCaller infers breakpoints both from the clipped and discordant alignments. We will describe the details of breakpoint identification later. On the other hand, if two normal pairs at both ends are removed after quality evaluation, then we will discard the whole candidate alignment since it is very likely the candidate alignment is a false alignment. Finally, each candidate alignment is evaluated by their numbers of exact matches and mismatches. For each short read, we only keep the alignment with the highest alignment score.

**Figure 1.**
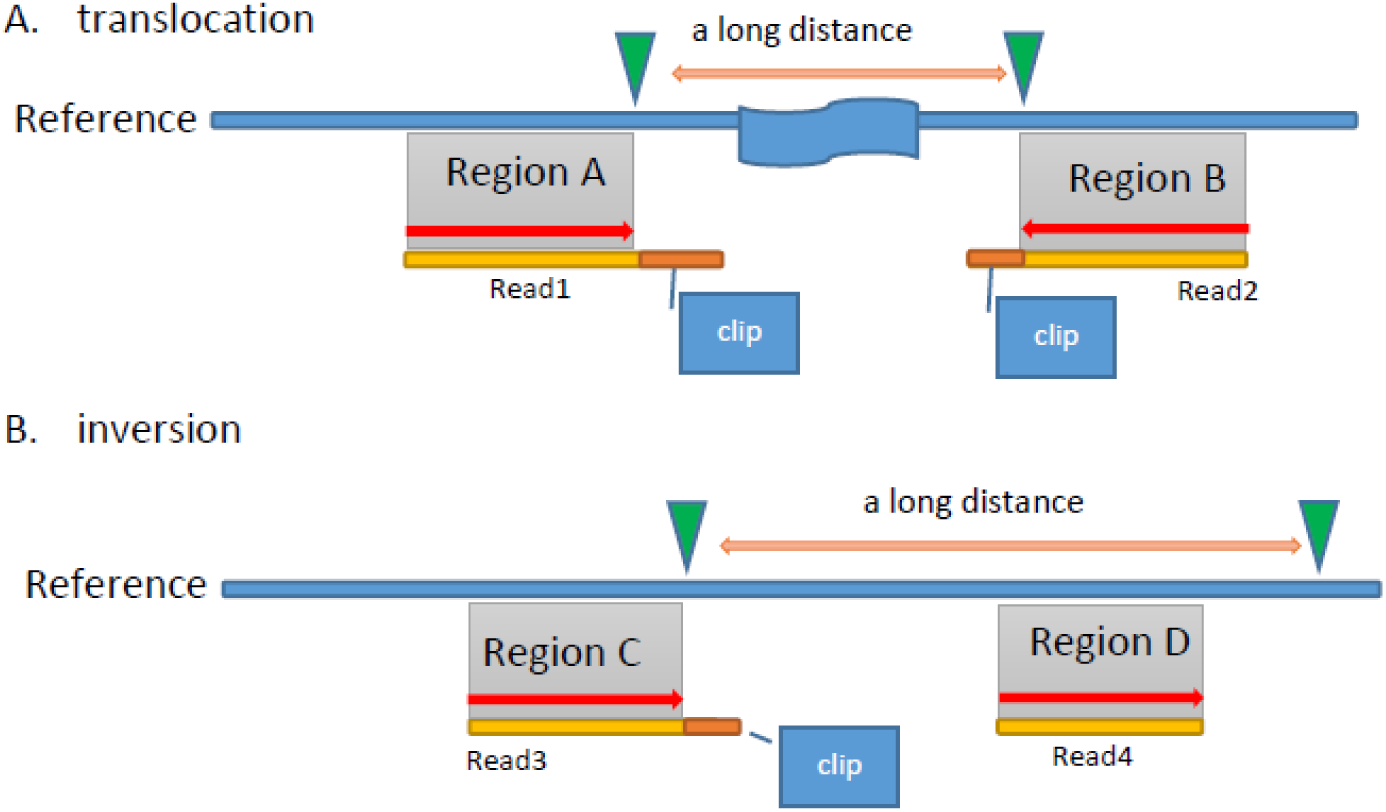
(A) Paired-end Read1 and Read2 are mapped distantly due to a translocation event in the sample genome. (B) Paired-end Read3 and Read4 are mapped with same orientation due to an inversion event in the sample genome.

MapCaller creates a position frequency matrix (PFM) to count the nucleobase occurrences at each position. PFM is a matrix of 4 × *L*, where *L* is the reference genome size. Therefore, each column of PFM represents the occurrences of nucleobases A, C, G and T at that position of the reference genome. MapCaller updates the occurrence at each column according to read alignments. Fig. 2 shows an example to illustrate how PFM works in this study. Five reads are mapped onto the reference genome. MapCaller counts the frequencies of each involved column. For example, PFM[3] = (1, 0, 0, 4) indicates there are one ‘A’ and four ‘T’s aligned at the third position of the reference genome. Any insertion and deletion events are kept otherwise. PFM can be further extended by increasing the column size if we would like to keep more mapping information. For example, we could add a field to count the read number starting at each position to filter out PCR duplicates, or we could also add a field to count the number of read orientation to avoid strand bias.

**Figure 2.**
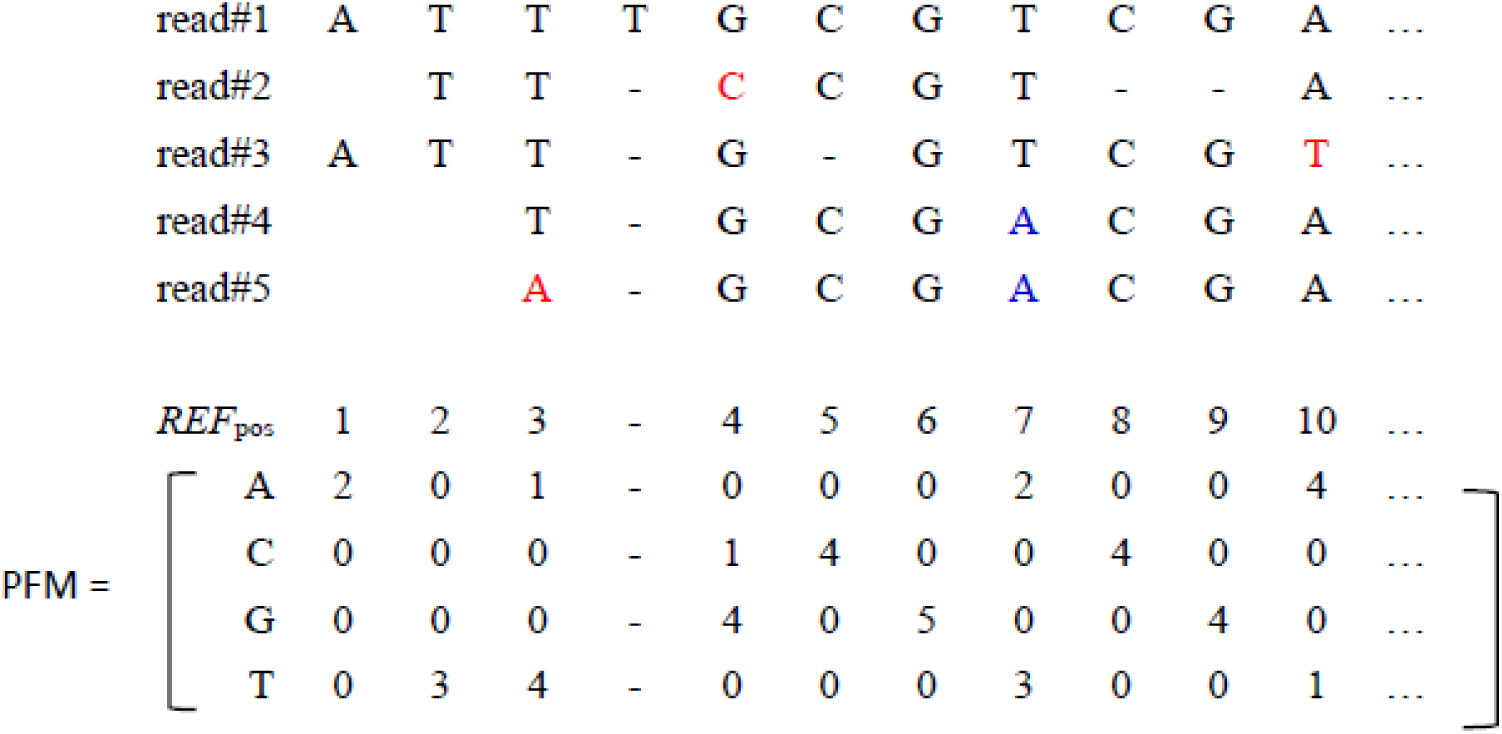
An example of position frequency matrix (PFM). Five read alignments are used to count the frequency of A, C. G and T at each position Nucleobases in red are sequencing errors, and those in blue are SNPs.

We use a 3-tuple, *Ins*(*Gpos, R*[*i,j*], *k*) and *Del*(*Gpos, G*[*m,n*], *k*) to represent an insertion and deletion event, where *Gpos* indicates the occurrence location, *R*[*i,j*] and *G*[*m,n*] are the indel strings, and *k* is the number of occurrences. MapCaller would create the following three 3-tuples: *Ins*(3, T, 1), *Del*(5, C, 1) and *Del*(8, CG, 1) based on the example cases in Fig 2. We describe how the SNVs, indels, inversions and translocations are identified based on the PFM and by MapCaller below. It is noteworthy that the quality measurement of each type of variant is described in the Supplementary (S1).

### SNV detection

MapCaller uses PFM to keep track of the occurrences of the four nucleobases at each position in the reference genome. The depth of position *p*, denoted as *Depth*(*p*), where *Depth*(*p*) = PFM[p, A]+ PFM[p, C]+ PFM[p, G] + PFM[p, T]. We partition the reference genome sequence into blocks of 100 nucleobases. For each block *i*, MapCaller determines a threshold, denoted as *depthr*(*i*), which is the half of the average depth for all the nucleobases within the block.

A nucleobase at position *p* is considered as an alternative allele if its occurrence is above *OccurrenceThr*(*p*) = *Depth(p)* × *MinAlleleFrequency*, where *MinAlleleFrequency* is a user defined threshold (the default value is 0.2 for germline mutation and 0.02 for somatic mutation). Thus, a nucleobase *b* at position *p* is considered as an SNV if the following two conditions are satisfied: (1) *Depth*(*p*) ≥ *depthr*(*i*); (2) PFM[p, b] ≥ *OccurrenceThr*(*p*). Since the sample genome may carry multiple sets of chromosomes, the genotype at the same position can be homozygous or heterozygous. MapCaller considers both haploid and diploid scenarios. If only one nucleobase is called and its frequency is less than 1-*MinAlleleFrequency*, then the locus is considered heterozygous, otherwise it is homozygous.

### Indel detection

MapCaller keeps track of all indel events using the 3-tuples, thus indel events can be deduced from those 3-tuples. We use *InsOccurrence*(*p*) and *DelOccurrence*(*p*) to represent the occurrences of insertion and deletion events at position *p*. However, the alignments involved with indels could be ambiguous. For example, the two following alignments produce identical alignment scores:

AGCATGCATTG

AGCAT----TG and AG----CATTG

It can be observed that the two alignments will lead to different indel events, which are GCAT and CATG at neighboring positions. To avoid the ambiguity, MapCaller finds the indel with the maximal occurrences within the range of (p −5, p + 5). MapCaller also counts the total occurrences of insertions and deletions (denoted as *InsOccurrence*(*p*) and *DelOccurrence*(*p*)) within the range. MapCaller reports an insertion event at position *p* only if *InsOccurrence*(*p*) ≥ *depthr*(*i*) × 0.25. Likewise, it reports a deletion event at position *p* only if *DelOccurrence*(*p*) ≥ *depthr*(*i*) × 0.35. Since *depthr(i)* is normally lower than the neighboring depths when there are deletion events, we use 0.35 to determine the threshold for deletion detection.

### Translocation and inversion detection

MapCaller estimates the average distance and fragment size of the paired-end reads periodically during read mapping. They are denoted as *AvgDist* and *AvgFragmentSize*. The definition of the distance and fragment size is described in Supplementary material (S2). We use *AvgDist* to distinguish concordant pairs from discordant pairs. In Illumina sequencing protocol, two paired-end reads are supposed to be mapped to different orientations of the same chromosome within an expected distance. Concordant pairs match paired-end expectations, whereas discordant pairs do not. Fig. 3(A) shows examples of regular paired-end reads that are mapped concordantly since the sample genome do not have any structural variants at the corresponding region. Fig. 3(B) shows a translocation event. The green dotted-rectangle is translocated from Breakpoint2 to Breakpoint1. Thus, the corresponding paired-end reads will be mapped discordantly. Likewise, Fig. 3(C) shows an example of an inversion event at BreakPoint3. The light-blue dotted-rectangle is the reversal of itself. Therefore, the corresponding paired-end reads will be mapped to the same orientation.

**Figure 3.**
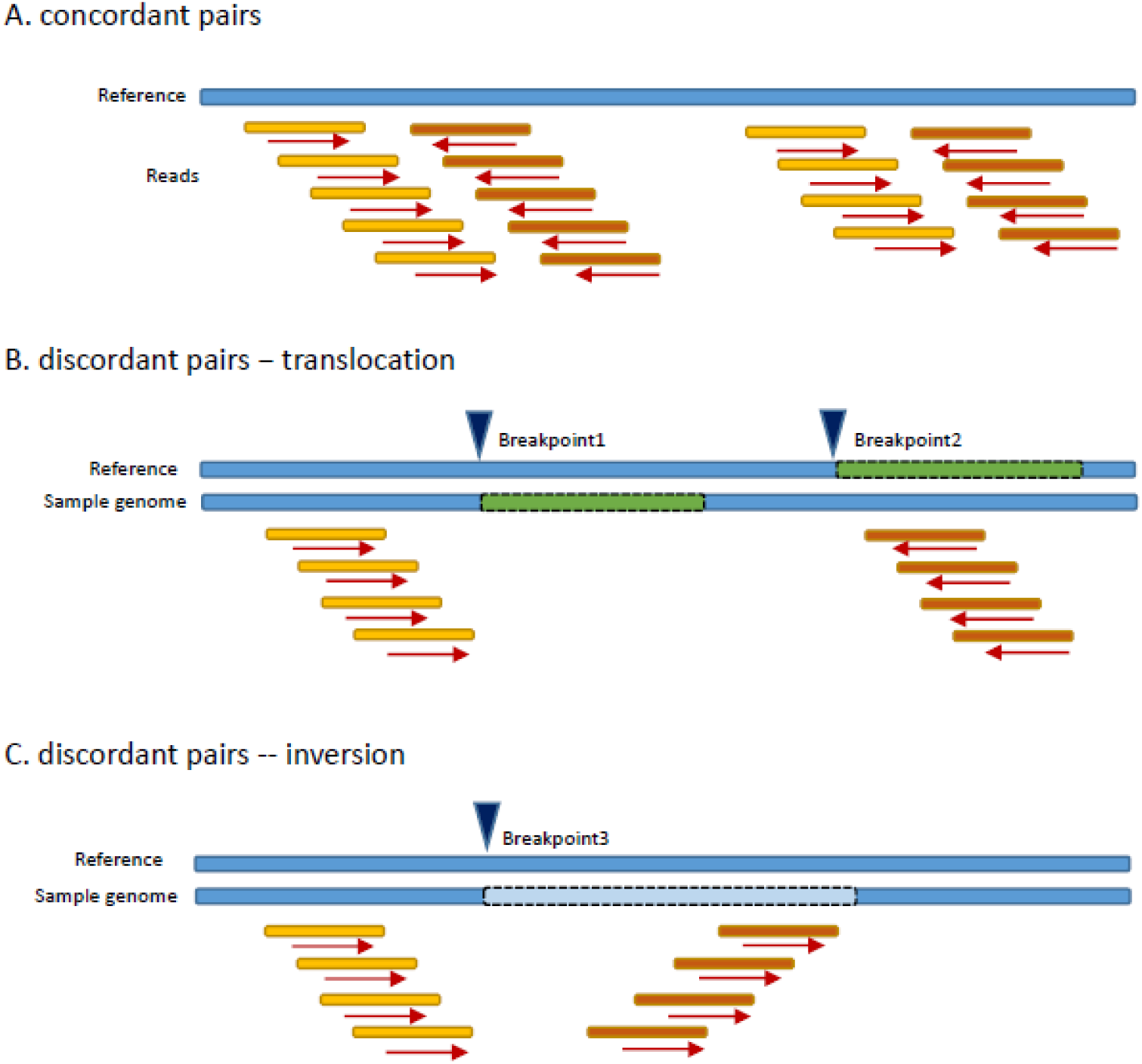
(A) Concordant pairs (B) discordant pairs-translocation (C) discordant pairs - inversion.

MapCaller collects all discordant pairs and classifies them into translocation and inversion candidate groups. If paired-end reads are mapped to different orientation and their distance is greater than (*AvgDist* × 1.5), they are put into translocation group, denoted as *TnlGroup*. If paired-end reads are mapped to the same orientation, they are put into inversion group, denoted as *InvGroup*. Since MapCaller collects all breakpoints from read alignments with a single-end clipping, we set a window to identify breakpoints as follows. For each breakpoint *p*, we set a screening region of *AvgFragmentSize* width at both sides of *p*. To identify breakpoints of translocation events, we count the number of reads in *TnlGroup* that are mapped at the screening region. The number of reads at left side is denoted as *LRnum*(*p*) and the right side is denoted as *RRnum*(*p*). We measure the average read depth at both sides, denoted as *Ldepth*(*p*) and *Rdepth*(*p*). If *p* is a true breakpoint for a translocation event, we should be able to observe a certain number of discordant read alignments in the *TnlGroup* that are separated by the breakpoint. Thus, we decide *p* is a breakpoint if the following conditions are satisfied: (1) *LRnum*(*p*) ≥ *depthr*(*i*); (2) *LRnum*(*p*) ≥ *Ldepth*(*p*) × 0.5; (3) *RRnum*(*p*) ≥ *depthr*(*i*); and (4) *RRnum*(*p*) ≥ *Rdepth*(*p*) × 0.5. To identify breakpoints of inversion events, we adopt similar process to verify the positivity of every breakpoint by checking the discordant pairs in the *InvGroup*.

## Data Availability

MapCaller was developed under Linux 64-bit environment and implemented with standard C/C++. It takes read files (FASTA/FASTQ) as input and outputs all predicted variants in VCF format. The source codes of MapCaller and benchmark datasets are available at https://github.com/hsinnan75/MapCaller.

## Acknowledgements

We would like to appreciate Prof. Torsten Seemann (Microbiological Diagnostic Unit) and Dr. Devon Ryan (Max Planck Institute of Immunobiology and Epigenetics) for their help with the software.

## Author contributions

H.N.L. conceived and designed the study; H.N.L. collected and pre-processed the datasets; H.N.L. developed the software; H.N.L. and W.L.H. wrote the manuscript. All authors read and approved the final manuscript.

