## Supplementary for "MapCaller – An integrated and efficient tool for short-read mapping and variant calling using high-throughput sequenced data"

Hsin-Nan Lin and Wen-Lian Hsu*
Institute of Information Science, Academia Sinica, Taipei, Taiwan

### Supplementary material

#### S1. The measurement of each type of variant in MapCaller

A. SNV at position p

QUAL = (35 × NS / Cov), where NS is the allele frequency and Cov is the read depth at *p*.

B. Indel at position p

QUAL = (100 × NS / Cov), where NS is the allele frequency and Cov is the read depth at *p*.

C. Translocation / Inversion at position p

QUAL = (-100 × log((1 - (NS / depthr(i))))), where NS is the frequency of the breakpoint *p*, and depthr*(i)* is the half of the average depth for all the nucleotides within the block *i*.

#### S2. The definition of paired-end read's distance and fragment size

Fig S1 illustrates the definitions of distance between paired-end reads and fragment size. Note that the distance is measured using only the coordination system of the forward strand. We estimate the average distance between paired-end reads to distinguish concordant pairs from discordant pairs. In Illumina sequencing protocol, two paired-end reads are supposed to be mapped to different strands of the same chromosome within an expected distance. Concordant pairs match paired-end expectations, whereas discordant pairs do not.

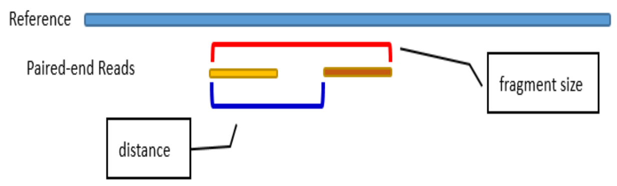


Fig S1. The distance and fragment size of two paired-end reads after mapping.

#### S3. The version and commands of MapCaller and selected existing tools

Table *S*1 lists the selected read mappers and variant callers tested in this study. The argument setting is also shown in the table.

Table S1. The version and commands of read mappers and variant callers

<table>
<tr><th>Tool</th><th>Version</th><th>Command</th></tr>
<tr><td>WGSIM</td><td>0.1.11</td><td>dwgsim -r 0 -C 30 -1 100 -2 100 <in.ref.fa> <out.prefix></td></tr>
<tr><td>Variant Simulator</td><td>-</td><td>var_sim <in.ref></td></tr>
<tr><td>MapCaller</td><td>0.9.9</td><td>MapCaller -t 12 -i ref_ix -f r1.fastq.gz -f2 r2.fastq.gz -vcf sample.vcf</td></tr>
<tr><td>KART</td><td>2.4.4</td><td>kart -t 12 -i ref_ix -f r1.fastq.gz -f2 r2.fastq.gz -o sample.sam</td></tr>
<tr><td>BWA-MEM</td><td>0.7.17</td><td>bwa mem -t 12 ref_idx r1.fastq.gz r2.fastq.gz > sample.sam</td></tr>
<tr><td>Bowtie</td><td>2.3.4.1</td><td>bowtie2 -x ref_idx -P 12 -1 r1.fastq.gz -2 r2.fastq.gz -S sample.sam</td></tr>
<tr><td>GEM</td><td>v3.6-7-gc2f9-dirty-release</td><td>gem-mapper -I ref_idx -t 12 -p -1 r1.fastq.gz -2 r2.fastq.gz -o sample.sam</td></tr>
<tr><td>Freebayes</td><td>v1.1.0-60-gc15b070</td><td>1. samtools view -@ 12 -b -h sample.sam > sample.bam<br>2. samtools sort -T sample -@ 12 -O bam sample.bam > sample.sorted.bam<br>3. Freebayes -f ref.fa sample.sorted.bam -v sample.vcf</td></tr>
<tr><td>SAMtools Mpileup</td><td>1.3.1</td><td>1. samtools view -@ 12 -b -h sample.sam > sample.bam<br>2. samtools sort -T sample -@ 12 -O bam sample.bam > sample.sorted.bam<br>3. bcftools mpileup -C 50 -Q 30 -Ou -f ref.fa sample.sorted.bam |bcftools call -vmO v -o sample.vcf</td></tr>
<tr><td>GATK</td><td>4.0.1.1</td><td>1. samtools view -@ 12 -b -h sample.sam > sample.bam<br>2. samtools sort -T sample -@ 12 -O bam sample.bam > sample.sorted.bam<br>3. java -jar picard.jar MarkDuplicates I=sample.sorted.bam O=sample.mark.bam M=sample.metrics.txt ASSUME_SORTED=true VALIDATION_STRINGENCY=LENIENT<br>4. java -jar picard.jar AddOrReplaceReadGroups I=sample.mark.bam O=sample.rg.bam RGID=1 RGLB=lib1 RGPL=illumina RGPU=unit1 RGSM=MyRead<br>5. java -jar picard.jar BuildBamIndex I=sample.rg.bam<br>6. gatk HaplotypeCaller -R ref.fa -I sample.rg.bam -O sample.vcf</td></tr>
<tr><td>DeepVariant</td><td></td><td>1. samtools view -@ 12 -b -h sample.sam > sample.bam<br>2. samtools sort -T sample -@ 12 -O bam sample.bam > sample.sorted.bam<br>3. docker run -v /exp_data:/input -v /exp_result:/output google/deepvariant:"0.9.0" /opt/deepvariant/bin/run_deepvariant /opt/deepvariant/bin/run_deepvariant --model_type=WGS --num_shards=12 --ref=ref.fa --reads=/input/sample.sorted.bam --output_vcf=/output/sample.vcf</td></tr>
<tr><td>Platypus</td><td></td><td>1. samtools view -@ 12 -b -h sample.sam > sample.bam<br>2. samtools sort -T sample -@ 12 -O bam sample.bam > sample.sorted.bam<br>3. Python bin/Platypus.py callVariants --nCPU 12 --bamFiles=sample.sorted.bam –refFile=ref.fa</td></tr>
</table>

| | | |
|---|---|---|
| | | –output=sample.vcf |
| DELLY | 0.7.7 | 1. samtools view -@ 12 -b -h sample.sam > sample.bam<br>2. samtools sort -T sample -@ 12 -O bam sample.bam > sample.sorted.bam<br>3. java -jar picard.jar AddOrReplaceReadGroups I=sample.sorted.bam O=sample.rg.bam RGID=1 RGLB=lib1 RGPL=illumina RGPU=unit1 RGSM=MyRead<br>4. delly call -t INV/BND -o sample.bcf -g ref.fa sample.rg.bam<br>5. bcftools view sample.bcf > sample.vcf |
| LUMPY | 0.2.13 | 1. samtools view -@ 12 -b -h sample.sam > sample.bam<br>2. samtools sort -T sample -@ 12 -O bam sample.bam > sample.sorted.bam<br>3. java -jar picard.jar AddOrReplaceReadGroups I=sample.sorted.bam O=sample.rg.bam RGID=1 RGLB=lib1 RGPL=illumina RGPU=unit1 RGSM=MyRead<br>4. samtools view -b -F 1294 sample.rg.bam > sample.disc.bam<br>5. samtools view -h sample.rg.bam \| scripts/extractSplitReads_BwaMem -i stdin \| samtools view -Sb - > sample.split.bam<br>6. lumpyexpress -B sample.rg.bam -S sample.split.bam -D sample.disc.bam -o sample.vcf |
| SVDetect | 1.3 | 1. BAM_preprocessingPairs.pl -p 1 sample.sam<br>2. ln -s sample.ab.sam mates/bwa.ecoli.ab.sam.all<br>3. SVDetect linking -conf sample.conf<br>4. SVDetect filtering -conf sample.conf<br>5. SVDetect links2SV -conf sample.conf |

#### S4. The variant simulator

We develop a simple program to modify a genome sequence to create simulated variants, including SNVs, indels, inversions and translocations. The source code of this simulator can be found at https://github.com/hsinnan75/MapCaller (folder: src/sv_simulator).

#### S5. Performance comparison result file

The detailed performance comparison result can be found at http://bioapp.iis.sinica.edu.tw/~arith/MapCaller/exp_result.xlsx
